# Temporally-precise basolateral amygdala activation is required for the formation of taste memories in gustatory cortex

**DOI:** 10.1101/2020.03.29.013995

**Authors:** Elor Arieli, Ron Gerbi, Mark Shein-Idelson, Anan Moran

## Abstract

Learning to associate malaise with the intake of novel food is critical for survival. Since food poisoning may take hours to affect, animals developed brain circuits to transform the current novel taste experience into a taste memory trace (TMT) and bridge this time lag. Ample studies showed that the basolateral amygdala (BLA), the nucleus basalis magnocellularis (NBM) and the gustatory cortex (GC) are involved in TMT formation and taste-malaise association. However, how dynamic activity across these brain regions during novel taste experience promotes the formation of these memories is currently unknown. We used the conditioned taste aversion (CTA) learning paradigm in combination with short-term optogenetics and electrophysiological recording in rats to test the hypothesis that temporally specific activation of BLA projection neurons is essential for TMT formation in the GC, and consequently CTA. We found that late-epoch (LE, >800ms), but not the early epoch (EE, 200-700ms), BLA activation during novel taste experience is essential for normal CTA, for early c-Fos expression in the GC (a marker of TMT formation) and for the subsequent changes in GC ensemble palatability coding. Interestingly, BLA activity was not required for intact taste identity or palatability perceptions. We further show that BLA-LE information is transmitted to GC through the BLA→NBM pathway where it affects the formation of taste memories. These results expose the dependence of long-term memory formation on specific temporal windows during sensory responses and the distributed circuits supporting this dependence.

**Significance:** Consumption of a novel taste may result in malaise and poses a threat to animals. Since the effects of poisoning appear only hours after consumption, animals must store the novel taste’s information in memory until they associate it with its value (nutritious or poisonous). Here we elucidate the neuronal activity patterns and circuits that support the processing and creation of novel-taste memories in rats. Our results show that specific patterns of temporal activation in the basolateral amygdala transmitted across brain areas are important for formation of taste memory and taste-malaise association. These findings may shed light on long-term activity-to-memory transformation in other sensory modalities.

## Introduction

A novel food poses a dilemma to animals: To eat or not to eat? On the one hand a new food may be highly nutritious, but on the other hand it may be toxic and life threatening (Rozin, 1976). To avoid the fatal consequences of poisoning, brain circuits have evolved to quickly detect novel tastes, transform these tastes into novel-taste memory trace (TMT)(Bermudez-Rattoni, 2014) to bridge the time-lag between consumption and malaise, and form a taste-malaise association (termed conditioned taste aversion [CTA]) (Garcia et al., 1955).

Decades of research have identified many brain regions that are involved in taste processing and memory. Among them, the basolateral amygdala (BLA), the gustatory cortex (GC) and the nucleus basalis magnocellularis (NBM) were acknowledged as key regions for taste novelty processing and taste memories acquisition (Bermudez-Rattoni, 2014; Yiannakas and Rosenblum, 2017). The GC is central to novelty processing, TMT formation and CTA learning (Bermudez-Rattoni, 2014; Moran and Katz, 2014; Yiannakas and Rosenblum, 2017). The novel-TMT formation in the GC is mediated by acetylcholine (ACh) secretion from NBM afferents (Gutierrez et al., 2003; Bermúdez-Rattoni, 2004). Correspondingly, ACh concentration in the GC is correlated with the novelty of the taste (Shimura et al., 1995; Miranda et al., 2000; Rodríguez-García et al., 2016). Other brain regions, however, likely activate the NBM to induce ACh secretion in the GC. A probable candidate is the BLA, which was implicated in novel taste processing. For example, BLA lesions or c-Fos blocking during CTA training reduces the CTA to a novel taste to the level of CTA to a familiar taste (Nachman and Ashe, 1974; Lamprecht and Dudai, 1996; Reilly and Bornovalova, 2005; St Andre and Reilly, 2007). In addition, BLA inhibition reduces taste neophobia – the diminished intake of a novel food relative to familiar food – implicating the BLA in taste novelty processing (Lin et al., 2018). Overall, these data suggest interactions between the BLA, NBM and GC underlie the processing of taste novelty.

Electrophysiological studies indicate that these interactions may be communicated through short time windows of neuronal activity. In the GC, taste stimuli elicit a sequence of short epochs of neuronal spiking activity containing chemosensory (early epoch [EE], 0.15-0.8 sec), palatability (late epoch [LE], 0.8-2 sec) (Katz et al., 2001; Fontanini et al., 2009; Maier and Katz, 2013) and novelty information (>2 sec) (Bahar et al., 2004). Interestingly, palatability is processed in the BLA during the EE (BLA-EE) (Fontanini et al., 2009) and inhibiting BLA during this epoch decreases palatability representation in the GC (Piette et al., 2012). This observation led to the hypothesis that palatability information “flows” from the BLA to the GC (Fontanini et al., 2009; Piette et al., 2012; Haley et al., 2016). Following the same logic, we hypothesize that BLA-LE activity is involved in novelty processing and that this information is transmitted to the GC, either directly (McDonald and Jackson, 1987; Haley et al., 2016), or indirectly through the NBM (McDonald and Jackson, 1987). The later seems appealing given the role of the BLA→NBM pathway in coding of arousing events (McGaugh, 2004).

To test these hypotheses we combined optogenetics, immunohistochemistry and electrophysiological recordings in behaving rats. Our results show that BLA-LE activity during novel-taste consumption is essential for the formation of CTA learning. Interestingly, this activity affects learning-related palatability coding changes in GC, without any influence on the perception of taste identity or palatability during training. In addition, our experiments confirmed the importance of the BLA→NBM pathway in CTA learning and showed that activating this pathway results in c-Fos expression in the GC, a marker of the novel-TMT formation. Together, our results suggest a new pathway by which specific BLA neuronal dynamics promotes the creation of taste memory in the GC.

## Materials and Methods

### Animals

Female Wistar rats (Harlan Biotech Israel Ltd) aged ~2.5 months and weighing 225-250g were used in this study. The rats were kept at the university animal facility under a 12h/12h light/dark routine, and while not under a specific experimental protocol received water and chow *ad libitum.* After having 2 days to acclimatize to their new home the rats were separated into individual cages. Rats were handled for 15 minutes a day for 3 days in order to habituate them for human touch. All methods comply with the Tel Aviv University Institutional Animal Care and Use Committee guidelines

### Surgeries

#### Anesthesia

Rats were temporarily anesthetized with isoflurane in an induction box, followed by an intraperitoneal injection of a ketamine-xylazine (KX) (100 and 10 mg/kg, respectively; maintenance: one-third induction dose every 1h).

#### Adeno-Associated Virus (AAV) injection

The anesthetized rat was placed in a stereotaxic frame, with its scalp excised, the scalp was opened and holes were bored above the BLA (AP = −2.9mm, ML = +4.85mm, measured from Bregma). Using an injector system (QSI, Stoelting) a syringe (Hamilton, 10μl, with 30G needle) was lowered slowly to depth of −7.6 from Bregma, and following additional 5 minutes to relieve tissue tension, was lowered to – 8.6mm. Following another 1 minute 0.8μl of viral vector (either experiment AAV_8_-CamKIIα-ArchT-GFP, or control AAV_8_-CamKIIα-GFP, University of North Carolina Vector Core) was injected in a flow rate of 0.1μl/minute (Gradinaru et al., 2008). At the end of the injection process the needle was left at the injection site for additional 5 minutes to allow virus spread. The needle was then pulled up another 1 millimeter, and following additional 2 minutes slowly withdrawn. The process was repeated on the other hemisphere. After the second injection, the rat was sutured and given post-operative care. Twenty one days were given for recovery and virus expression prior to fiber-optic and electrode implantation.

#### Fiber-optic and electrode implantation

The anesthetized rat was placed in a stereotactic frame and the scalp was reopened. The skull was thoroughly cleaned using sterile que tips, Kimwipes and bleached with peroxide (30%). Four self-tapping ground screws were inserted into the skull. The previously-drilled holes above the BLA were cleaned to allow the insertion of two optic fibers (200μ, 0.39NA, FT200UMT, Thorlabs) with metal ferules (Thorlabs, 2.5mm) that were implanted above the BLA (AP = −2.9mm, ML = ±4.85mm, DV = −8 mm, from Bregma) or NBM (AP= −1.4mm, ML=±2.8mm, DV=−7, from Bregma). For electrophysiological experiments an additional hole was drilled above the GC (AP = 1.4mm, ML = 5mm, DV = −4.5mm, relative to Bregma), and the dura was removed. Either one or two self-made movable 32 electrode bundles (Piette et al., 2012; Moran and Katz, 2014) were then lowered into the GC, and secured to the skull with dental acrylic. In addition, two intraoral cannulas (IOC, flexible plastic tubing, AM-Systems) were inserted through the oral cavity lateral to the second molar tooth (Phillips and Norgren, 1970; Piette et al., 2012; Moran and Katz, 2014). When finished the entire structure was covered with dental acrylic. Rats were put in individual cages and given at least 7 days to recover from the second surgery.

#### Optrode implantation and recording

We used optrode to confirm our ability to inhibit the BLA projection neurons in prolonged and short stimulation. The optrode was made by gluing 8 electrodes to same optic fiber used in our optogenetic experiments and cutting them ~0.5 mm below the end of the fiber. The optrodes were implanted similarly to the optic fiber implantation. Following 7 days of recovery, the rats were connected to the recording and optogenetic systems and neuronal activity from the BLA neurons was recorded with and without the laser pulses of 3 and 0.5 seconds. Laser power was set to achieve 30-40mW illumination at fiber tip.

#### Post-operative treatment

Following surgery, the rats were given subcutaneous injections of antibiotics (5mg/kg of Baytril 5%), pain relievers (1mg/kg of Meloxicam 0.5%) and Saline (10ml/kg) to ensure hydration. The head wound margins were treated with antibiotic cream.

### Experimental Design

#### CTA paradigm

Following recovery, water was removed from the home-cage. The basic experimental procedure is depicted in Figure 2A, B. Over the initial 3 experiment day days the rats were habituated to poke in an infrared-operated nose-poke (Coulbourn Instruments) for 40μl drops of water delivered through the IOC in the morning session (with 3 seconds inter-stimulus interval [ISI]), and to 30 minutes access to a water bottle several hours later (to ensure proper hydration). On day 4 the rats were given a 0.2M sucrose solution instead of water as a stimulant. Importantly, sucrose was chosen as a stimulant as sucrose evokes only minimal neophobic reaction (Miller and Holzman, 1981), thus exclude possible impact of BLA optogenetic inhibition over neophobic process. During sucrose IOC deliveries the BLAs of the rats were inhibited in different epochs according to the specific experimental group (Fig. 2B, green bars). Taste delivery times initiated by crossing the infrared beam in the nose poke were recorded by the electrophysiological acquisition system (Inten Technologies). Following 20 minutes the session ended, the rats were returned to their home cage, and twenty minutes later were given an intraperitoneal injection of 0.3M Lithium Chloride (LiCl, 1% body weight) to induce malaise. The next morning (day 5) the rats were placed again in the experimental chamber and were offered to poke for sucrose as in day 4. The reduction in their willingness to poke was used to measure the effects of the CTA learning (presented as % of pre-CTA pokes).

#### *CTA paradigm* with electrophysiology

The experimental procedure involving electrophysiological recording in the GC is shown in figure 6B. The procedure mimics the previously described CTA paradigm, but here the free-poking session was followed by a ~40 minute passive delivery session in which drops of tastes were delivered through the IOC to record neuronal taste responses (with a 20 seconds ISI). During the first 3 habituation days water, citric acid (CA, 0.1M) and salty (NaCl, 0.1M) solutions were offered through the IOC, and sucrose (Suc, 0.2M) was added to these 3 tastes in the 4^th^ (Pre-CTA) and 5^th^ (Post-CTA) days.

#### BLA optogenetic inhibition (BLA_ox_) paradigm

A 532nm green laser (CNI, 150mW) was connected to each of the two fiber optics ferrules and controlled using a TTL signal from an Arduino board. Laser strength was set according to the specific implanted fiber characteristics to achieve 30-40mW at the tip. Laser onset and duration were set according to the specific experimental group and coupled to taste delivery. The different epochs used were (in milliseconds from taste delivery): 0-3000 (Full), 0-500 (EE), 700-1200 (LE), 2500-3000. In one group we extended the ISI to 5000 and used 3500-4000 BLA_ox_.

#### State-dependency and double CTA procedure

Rats were subjected to the same experimental paradigm as described above in “CTA paradigm”. However, the rats that tested state-dependency received BLA_ox_ during the testing session in addition to the one in the training session (Fig. 2B, “Pre+Post1”). Following the test session these rats received a second LiCl injection and were tested a second time the day after to test their ability to acquire CTA with multiple CTAs (Fig. 2B, “Pre+Post1+post2”).

#### Go/No-Go paradigm

Rats were trained to poke for water as described previously in “CTA paradigm”. Following habituation to the poking paradigm they were introduced to a new paradigm in which pseudo-random number of water trials (3-7) were followed by a drop of 0.2M sucrose solution. Rats were required to identify the sucrose and withdraw from the nose poke for at least 5 seconds. While a successful withdrawal restarted the water trials, a failure to exit or remain outside long enough was punished by an aversive 10mM quinine drop. After the rats reached a plateau of at least 70% correct trials we started daily sessions in which we performed full BLA_ox_ simultaneously with sucrose deliveries in random half of the trials. To eliminate light information from guiding the rats’ decision an additional non-implanted optic fiber was turned on during all the trials. BLA_ox_ impact on sucrose identification was assessed by comparing performance between BLA_ox_ and non-BLA_ox_ within the same animal.

### Palatability assessment based on poking microstructure

Previous studies have shown that palatability can be inferred from the length of lick bouts (Davis and Perez, 1993; Hsiao and Fan, 1993). Our use of poking instead of licking required modification of this method (graphically presented in figure 3D, top). Poking times collected by our system in response to nose pokes were stored offline. We defined a poking bout as 5 consecutive pokes or more without withdrawal from the nose poke for more than 10 seconds. We then used these data in order to calculate average bout length (measured as pokes count) and the number of bouts. Since the 3500-4000ms BLA_ox_ group used a longer minimum ISI (5 seconds) it could not be compared with the other groups and therefore was not analyzed.

### Perfusion

The rats were first anesthetized with KX solution and then perfused with saline (0.9% NaCl) followed by 4% formaldehyde solution. After fixation the brain was extracted from the skull and left for 72 hours in a 30% sucrose formaldehyde solution at 4°C until sliced and mounted on slides.

### Histology

The fixed brains were cut to 50μm slices using a microtome (Fisher Scientific), plated on microscope slides, covered with DAPI containing preservative (<u>Invitrogen Flouromount-G with DAPI</u>) and left to dry for 24 hours. Spread and expression of the virus and the correct location of the optic fibers above the BLA were performed using fluorescent microscope. Only rats with bilateral correct fiber location and sufficient virus expression were included in the study.

### Novel taste paradigm for c-Fos expression

Rats that were used in the c-Fos experiments followed the same AAV infection, fibers and IOC surgeries as previously described for the behavioral experiments. The control groups were infected with sham virus and implanted with fibers and IOC similar to the experimental group. Following at least 7 days of recovery the rats were habituated to a watering schedule for 3 days as previously described. On the 4^th^ day the different groups received sucrose solution (or water, in the water groups) with or without BLA_ox_ during either full (0-3000) or LE (700-1200) ms post taste delivery. Importantly, the control groups received laser stimulation but without a functional virus. At the termination of the experimental session the rats were returned to their home cage. Exactly 90 min afterwards the rats were anesthetized with KX (100mg/10kg, injected IP), perfused transcardially with 0.1M phosphate-buffered saline (PBS, pH 7.4) followed by 4% paraformaldehyde (PFA) in PBS. Afterwards, brains were removed and post-fixed in 4% PFA for 24 hr, followed by 30% sucrose solution in PBS for 72hr at 4°C. Coronal sections of 50 μm were cut using a microtome (Fisher Scientific) and were collected into a cryoprotectant solution (25% Glycerol and 25% Ethylene Glycol in 0.1 M PO_4_) and stored in 4°C until immunofluorescence assay.

### Immunofluorescence assay

Free-floating sections were rinsed (3 × 10 min) with PBS, then permeabilized for 4h in PBS containing 0.3% Triton X-100 (PBS-Tx), followed by 3h blocking with 2.5% Normal Goat Serum in PBS-Tx at room temperature. Sections were then incubated with the rabbit anti-Fos primary antibody, 1:1000 (Cell Signaling, 2250S) in block at 4°C overnight. Following, sections were rinsed (3 × 10 min) with PBS, incubated with the secondary antibody Alexa Fluor^®^ 594 Conjugate goat anti rabbit, 1:1000 (Cell Signaling, 8889S) and with DAPI 1:1000 in block for 3h at room temperature. Sections were rinsed 3 × 10 min with PBS, mounted on slides, and cover slipped with mounting medium.

### Image Acquisition and Quantification

Images were acquired using an Olympus epifluorescence microscope (10 × 0.45 NA or 20 × 0.75 NA objective). Quantification of c-Fos positive nuclei was based on 2 images of the GC region (Bregma 1.5 – 0.0, Paxinos & Watson, 1998) per hemisphere per rat. The threshold was calculated for each image separately (Yasoshima et al., 2006). Each image was processed to reach a binary image using the ImageJ software (NIH), followed by automated quantification of c-Fos positive nuclei that overlaid with DAPI staining to ensure localization in the nucleus. Counts were averaged per rat and over group.

### Acquisition and analysis of electrophysiological data

Acquisition and pre-processing: Extracellular neuronal signals were collected from self-manufactured (Piette et al., 2012; Moran and Katz, 2014) 32-wire electrodes (0.0015” formvar-coated Nichrome wire; AM Systems) positioned within the GC, during taste deliveries. The data was first collected by an Analog-to-Digital head-stage amplifier (RHD2132, Intan Technologies) and then sampled at 30Khz by an Intan RHD2000 acquisition system and stored offline. Common noise was removed from each recorded channel using a common average reference (CAR) algorithm. Spike sorting was performed using the opensource KlustaKwik/Phy software (Rossant et al., 2016). Initially, putative spikes were detected using a 3-6 standard deviation threshold, and then automatically clustered. Using the Phy software the clusters were further manually classified as noise, multiunit or single unit spiking activity based on the spike shape, apparent refractory period in the autocorrelation plot and separation from the noise cluster. Only well isolated single neurons were added to the cohort used in study (Piette et al., 2012; Moran and Katz, 2014), and less than 10% of the electrodes recorded more than a single unit.

### Single-unit palatability distance ratio calculation

To assess whether sucrose palatability in a certain neuron is coded as aversive or palatable we compared its sucrose response firing rate to responses to aversive citric acid or palatable salty solution. Specifically, we defined the response distance (RD) as the difference between the firing rate of a neuron in the 700-1200ms window to sucrose and either aversive 0.1M citric acid (D[S~C]) or palatable NaCl 0.1M (D[S~N]). RD was calculated as the Euclidean distance between two N-dimensional vectors containing the neuron’s firing rate during N 50ms time bins (see graphical representation of RD calculation in Fig. 7A). We then calculated for each neuron the palatability distances ratio as 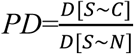. The higher the PD (above 1), the more sucrose is coded as palatable.

### Neuronal population palatability evaluation

NaCl, CA and sucrose trials from the forced taste delivery part of the CTA paradigm (second part of each daily session, Fig. 6B) were used for the classification of sucrose trials. Taste neuronal responses were averaged over the bins corresponding to the epochs tested (0-250ms, 250-500ms, 500-750ms, 750-1000ms, 1000-1250ms, 1250-1500ms) for each neuron in order to receive an N-dimensional population response vector for each trial in each epoch (N = collective number of neurons from an experimental group). We then used random 70% of the NaCl and CA trials to train a Multi-Layered Perceptron taken from the Scikit-learn package with 2 hidden layers of 100 and 50 cells. Following training, the performance of the network was tested with the remaining 30% trials of NaCl and CA responses and the amount of correctly predicted trials was calculated. Sucrose trials were fed into the same network and the probability of sucrose being identified as NaCl was received. This entire process was repeated 200 times for each epoch in each group and averaged out in order to eliminate noise based on starting conditions or the selection of a specific train/test split.

### Statistical analysis

Statistical analyses were conducted using Python, specifically the statsmodels and scipy packages, and SPSS (IBM). ANOVAs followed by post-hoc t-tests were conducted to evaluate the statistical significance of the differences between groups or experimental procedures when the data was normally distributed. When the data was not normally distributed Kruskal-Wallis was used followed by post-hoc Mann–Whitney U tests. Two-tailed tests were used unless otherwise stated. The significance level was set at α = 0.05. Data statistics, including all error bars, are presented as mean ± standard error of the mean unless otherwise stated.

## Results

To study the importance of short-term BLA projection neuron activity in taste perception and learning we employed an optogenetic approach. Specifically, bilateral BLA infection with AAV_8_-CamKIIα-ArchT-GFP (AAV_8_-CamKIIα-GFP in control groups) targeting the excitatory projection neurons (Fig. 1A) was followed by bilateral fiber optic implantation with the tips located within and above the BLA (Fig. 1B). Post-experiment analysis of brain slices confirmed wide-spread infection within the BLAs (Fig. 1C), as well as the correct localization of the fiber tips (Fig. 1D). Only rats that exhibited both virus expression and localization, and correct fiber positioning were included in this study. We confirmed our ability to optogenetically inhibit the BLA for short (0.5 sec) and long (3 sec) timespans with limited rebound excitability (Madisen et al., 2012; Mattis et al., 2012). This was done using an optrode device (Fig. 1E, see the Methods section) capable of simultaneous recording and optogenetic manipulation within the same brain region. Example recordings from the BLA with long (3 seconds) and short (0.5 sec) BLA_ox_ are presented in Fig. 1F and Fig. 1G, respectively. These recordings confirmed that under our BLA_ox_ protocol, neurons exhibit fast inhibition, and quickly return to normal firing when the light was turned off, with a minimal rebound excitation. With this confirmation in hand, we proceeded to test the roles of the BLA in taste perception, memory and learning.

**Figure 1:**
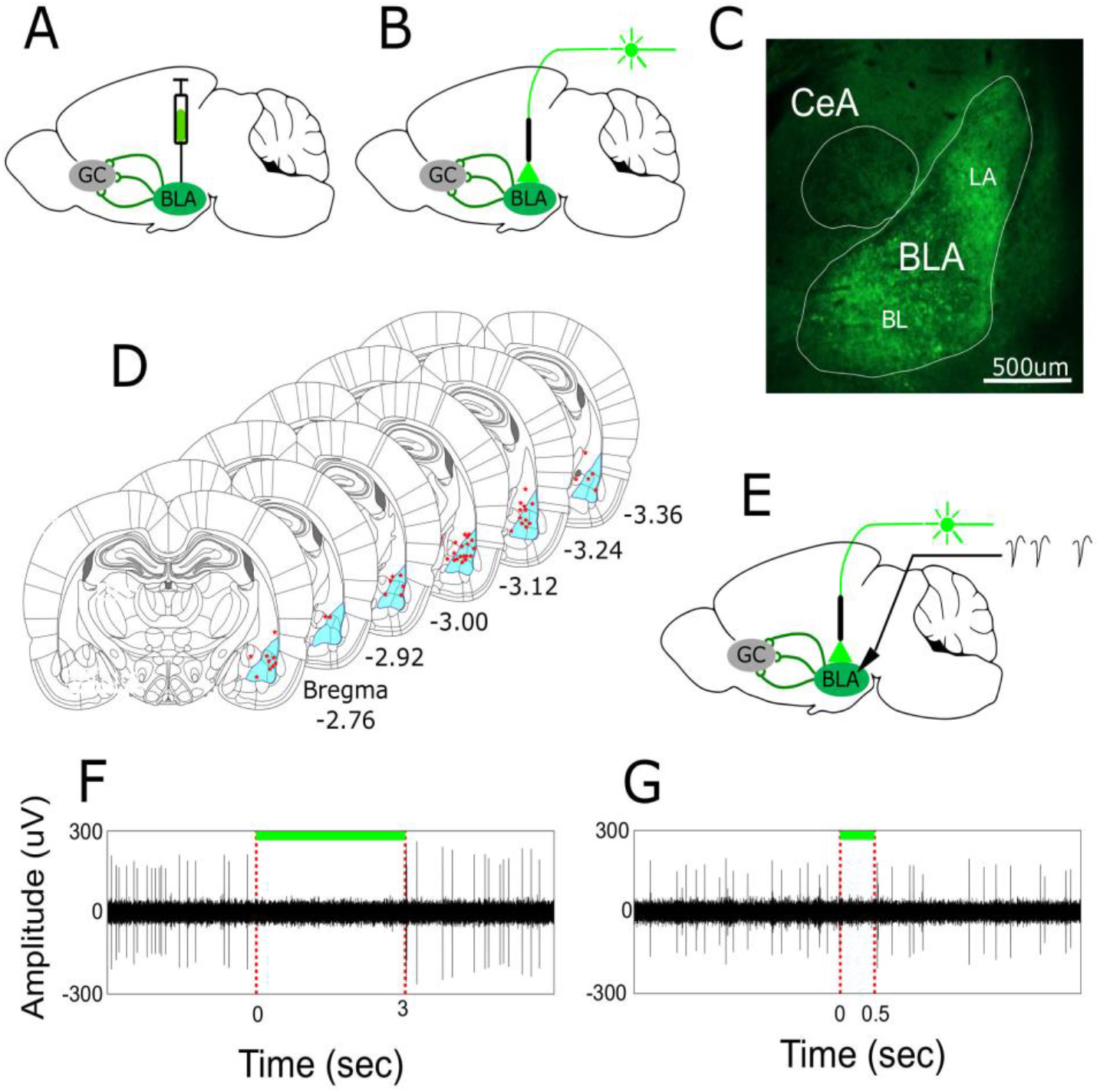
Verification of viral expression and optogenetic inhibition of the BLA. A) Rats were first infected with AAV_8_-CamKII-ArchT-GFP (AAV_8_-CamKII-GFP for controls) viral vector injected to the BLA, bilaterally. B) In a second surgery 21 days after infection, fiber optics were implanted above the BLAs, together with bilateral IOCs for taste deliveries C) A coronal section from a rat infected with AAV_8_-CamKII-ArchT-GFP construct, showing ArchT expression (tagged by GFP) localized in the BLA complex. Notice that the central amygdala (CeA), which mostly contains inhibitory GABAergic neurons not targeted by the virus, is visible but lack the GFP staining. D) Localization of fiber-optic tips in the rats used in this study. The tip locations from both hemispheres are collapsed on a single hemisphere for better visibility. E) Schematics of the BLA inhibition verification using an optrode device implanted in the BLA. F-G) Example of BLA neuronal activity responses to laser illumination recorded by the optrode in a behaving rat. Both long 3 seconds (F) or short 0.5 second (G) BLA_ox_ caused immediate cessation of neuronal activity followed by a quick return to normal firing after offset.

### Prolonged optogenetic BLA inactivation during novel taste experience attenuates CTA learning

Our first aim was to confirm our ability to attenuate CTA acquisition for a novel taste using prolonged BLA optogenetic inhibition, as was previously reported following BLA lesion (Nachman and Ashe, 1974; Rollins et al., 2001; Reilly and Bornovalova, 2005; St Andre and Reilly, 2007) and pharmacological interventions (Lamprecht and Dudai, 1996; Berman et al., 1998, 2000; Koh and Bernstein, 2003). To that end, we trained mildly water-deprived rats to poke for drops of water delivered through the IOC (Fig. 2A, B, see the Materials and Methods section). This technique combines clear assessment of behavior with precise control over stimulus timing. On the training day, sucrose replaced water as a stimulant and the session was repeated, but now every drop of sucrose was accompanied by BLA_ox_ for the 3 seconds ISI, effectively blocking the BLA for the entire tasting period. To control for possible heat related effects of the optogenetic stimulation, the control group received similar laser stimulation as the experimental group but lacked a functional ArchT channel in the BLA. Following this session, the rats were injected with the emetic LiCl to induce malaise, and consequently CTA. The strength of the CTA was tested the day after when the rats were offered to poke again for sucrose (Pre-only group).

**Figure 2:**
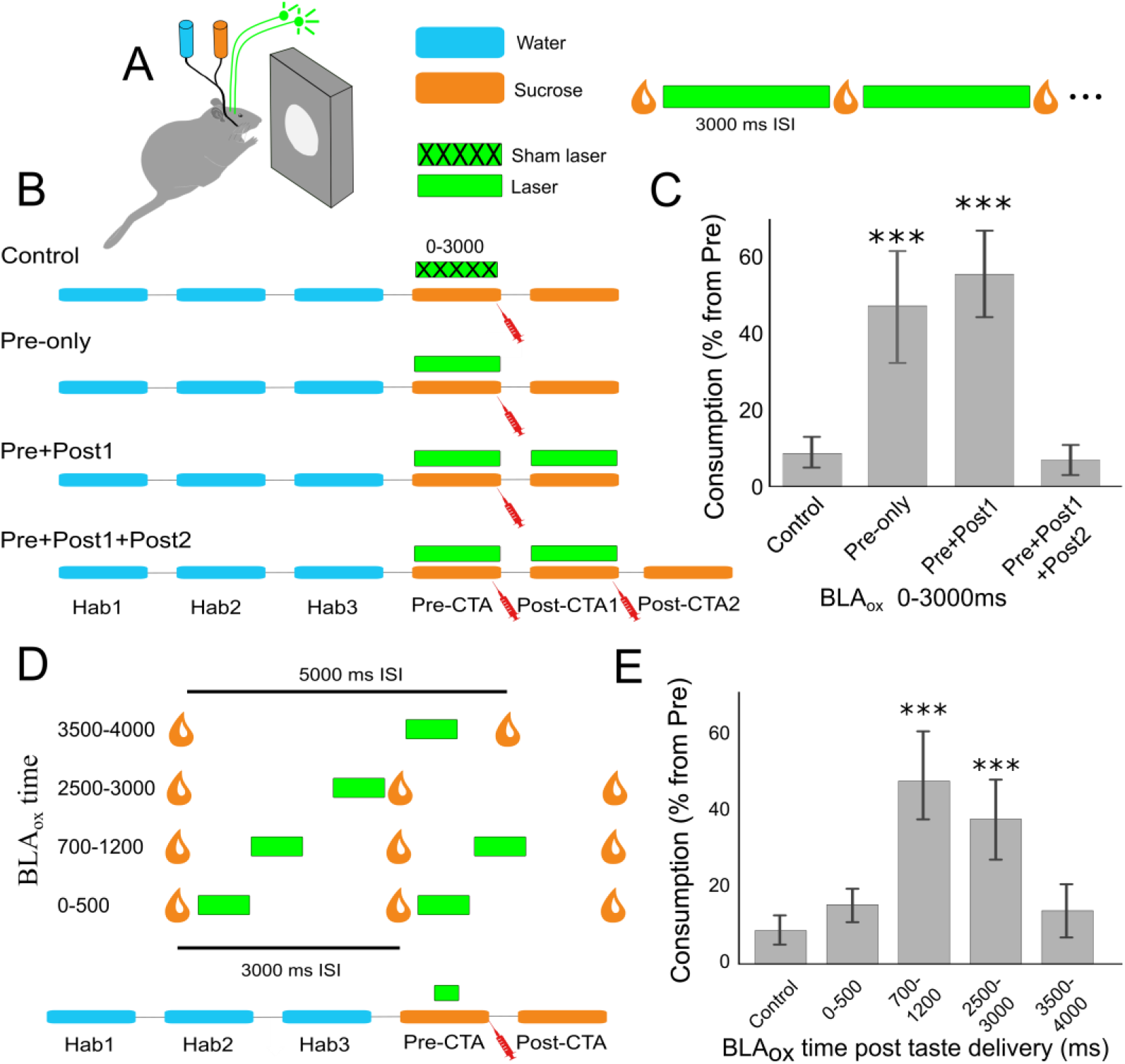
Short-term BLA activity is required for CTA acquisition. A) Schematics of the experimental design. Rats were implanted with two optic fibers in their BLAs and an IOC that delivers drops of liquid directly into the oral cavity in response to a nose-poke. Right: This experimental system provided precisely-times BLA_ox_ with respect to taste deliveries. B) Prolonged BLA_ox_ experimental protocol. During the initial 3 habituation days (Hab) the rats were trained to poke for drops of water (W) which were continuously delivered with an ISI of 3000ms. On the Pre-CTA day, water was replaced with 0.2M sucrose (Suc) and BLA_ox_ was performed over the entire 3000 ISI. This session was followed by a LiCl injection to induce malaise (red syringe). On the following day, rats were offered to poke for sucrose drops and the total number of drops (Consumption) relative to Pre-CTA (% from Pre) was measured to access aversion level. Control and Pre-only groups received the same protocol. The Pre+Post1 group received BLA_ox_ also during the first Post-CTA day. Pre+Post1+Post2 group received additional LiCl injection after the first Post-CTA day, and tested again a day later. C) Suc consumption for the groups in B (B). Control group showed strong aversion to sucrose. Pre-only (n=6) and Pre+Post1 (n=3) groups showed significant attenuation of CTA. Additional LiCl injection following testing (Pre+Post1+Post2, n=3) caused similar rejection of sucrose as the control rats, confirming CTA learning. D) Short-term BLA_ox_ experimental protocol. Different groups received 500ms BLA_ox_ at specific epoch following each taste delivery in the Pre-CTA day. E) Consumption for the experiments in (D). BLA_ox_ performed during the late epoch (700-3000ms) shows attenuated aversive memory formation. *** p<0.001 tested vs. the control group. Control n=6; 0-500 n=5; 700-1200 n=10; 2500-3000 n=6; 3500-4000 n=7.

The results of this experiment clearly demonstrate the attenuation of CTA memory acquisition following BLA inhibition: the BLA_ox_ Pre-only group showed significantly reduced CTA learning compared with the control group (Fig. 2C; t-test, p=6.5×10^-5^). We further confirmed that the attenuation of learning is not the result of the laser becoming part of the conditioned stimulus (a generalization decrement effect (Capaldi, 1994; Bouton, 2004)) and therefore causing a weaker response in the testing day when the laser is absent. To that end, we added a group in which rats received BLA_ox_ during both the training and testing days (Pre+Post1 group, Fig. 2B). This group showed attenuation of the CTA learning compared with the control group (t-test, p=3×10^-5^) which was similar to the BLA_ox_ Pre-only group (Fig. 2C, t-test, p=0.222), and therefore rejects a generalization decrement (state dependency) effect. An additional injection of LiCl given to the Pre+Post1 group at the end of the first testing session (Fig. 2B, Pre+Post1+Post2) induced normal CTA (Fig. 2C, t-test vs. control p=0.624), indicating that BLA_ox_ attenuates, but does not block the CTA, as was previously reported in BLA lesion studies (Reilly and Bornovalova, 2005; St Andre and Reilly, 2007). Taken together, these results confirm that perturbing the initial tasting experience interferes with the creation of a long-term CTA memory.

### CTA learning requires the activation of BLA projection neurons during specific epochs

Previous studies reported that neuronal activity in short epochs during GC and BLA taste responses are correlated with taste qualities such as identity, palatability and novelty (Katz et al., 2001; Bahar et al., 2004; Jones et al., 2007; Fontanini et al., 2009; Maier and Katz, 2013; Moran and Katz, 2014). We thus went on to test whether BLA activity during a specific epoch carries information that is important for CTA formation. To that end, we repeated the CTA procedure with BLA_ox_ during training as previously described, but now different groups of rats were subjected to BLA_ox_ only during a specific short (500ms) epoch following each drop of taste delivery (Fig. 2D). Interestingly, we found that the groups differ in their ability to acquire the CTA memory a day later (Fig. 2E, One-way ANOVA, F_(4,32)_=11.75, p<0.001). While the group receiving BLA_ox_ during the early epoch (0-500ms, EE-BLA_ox_) showed normal sucrose avoidance following the CTA, short BLA_ox_ during the late epoch (>700 and <3000ms) significantly attenuated CTA memory formation (Fig. 2E; t-test vs. control; p<10^-5^, p<10^-4^ for 700-1200ms and 2500-3000ms groups respectively). This memory attenuation was similar to that of the BLA_ox_ group inhibited for the entire 3000ms (Fig. 2E; t-test vs. Pre-only group, p=0.992, p=0.358 for 700-1200ms and 2500-3000ms groups respectively). By extending the ISI to 5000ms we found that the late boundary of the LE is around 3500ms since BLA_ox_ during 3500-4000ms resulted in normal CTA learning (Fig. 2E). These results show that LE activity in BLA projecting neurons during novel taste experience, but not the EE activity, is required for intact CTA learning.

### Activity of BLA projection neurons is not required for taste identity and palatability perceptions

What might be the role of the BLA during novel taste experience, and specifically during the LE, that gives rise to the attenuation of CTA? One possibility is that it interrupts with the perception of taste identity. Although the BLA is not considered a part of the main taste system, BLA_ox_ might still interrupt the processing of taste identity in other downstream brain regions. To rule this out, we trained rats in a Go/No Go task. Rats learned to poke for drops of water delivered through the IOC as before, but withdraw once they identify a drop of sucrose in order to avoid the delivery of a highly bitter quinine solution 3 seconds later (Fig. 3A, see also the Material and Methods section). The rats improved their performance over days and reached at least 70% correct performance in 4 days (Fig. 4B). Following reaching a stable performance, 0-3000ms BLA_ox_ was performed during random sucrose presentations in half of the trials within the same session. We found similar performance between the BLA_ox_ and non-BLA_ox_ trials across 4 test days (Fig. 3B, Two-way ANOVA, Day: F_(1,2)_=0.11, p=0.89; BLA_ox_ F_(1,2)_=0.961, p=0.34, interaction: F_(1,2)_=2.253, p=0.13). As expected, our results suggest that misperceived taste identity due to BLA inhibition cannot account for the attenuation of CTA by BLA_ox_.

**Figure 3:**
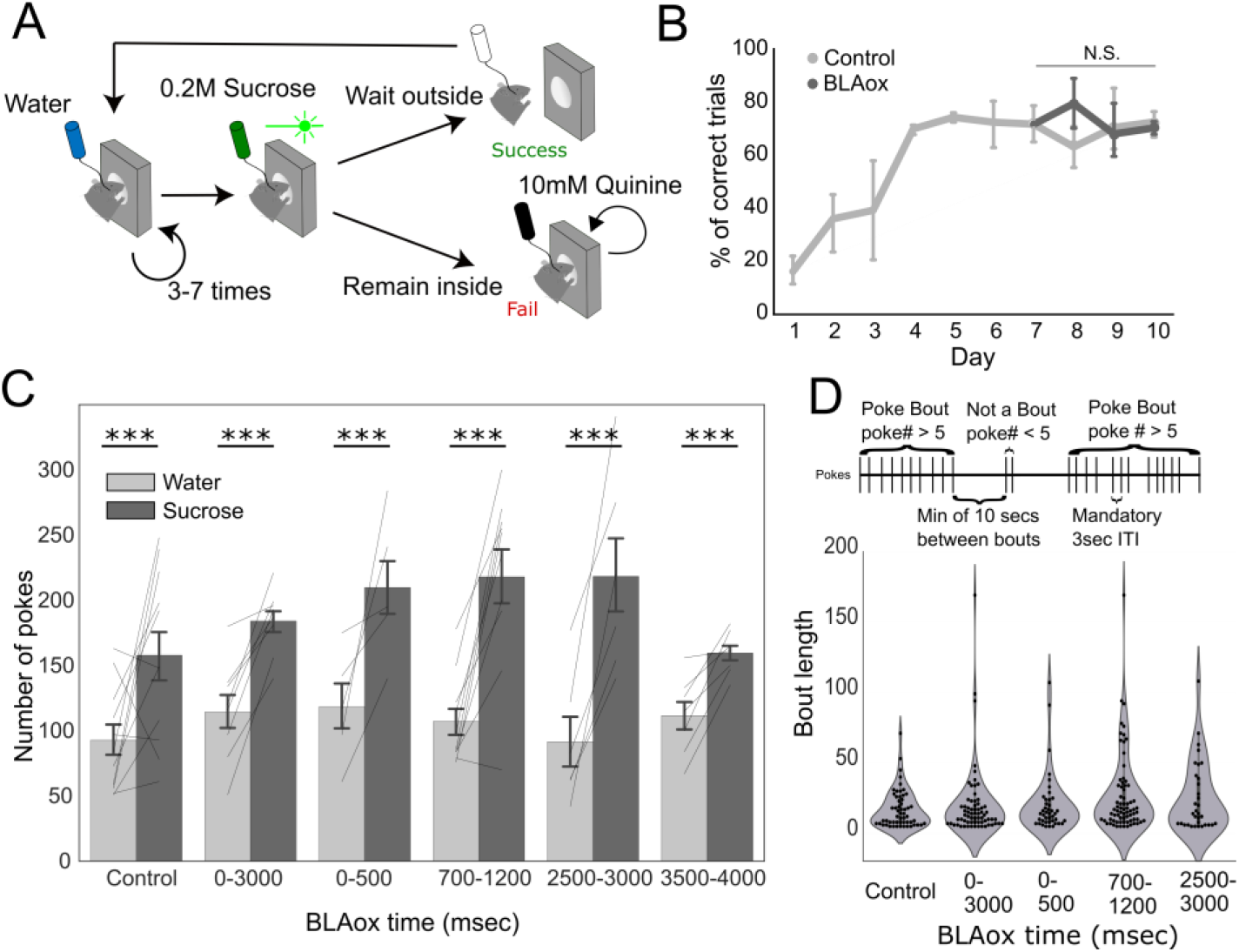
Effects of long-term BLA_ox_ on taste perception and learning. A) A schematic of the Go/No-Go experimental paradigm. Rats were trained to poke for random number (3-7) of water deliveries before receiving a single drop of sucrose. The rats were required to identify the sucrose taste and withdraw from the nose poke for at least 5000ms in order to avoid a highly bitter 10mM quinine droplet. Following reaching a plateau performance for 3 consecutive days, half (randomly chosen) of the sucrose deliveries were performed with full BLA_ox_. B) Percentage of correctly identified taste trials across days. BLA_ox_ showed no significant effect on the ability of the rats to identify the sucrose taste. C) The number pokes for water (in the last habituation day) and for 0.2M sucrose (in the training day) prior to CTA. All BLA_ox_ groups poked significantly more for sucrose than water, similar to the control group, showing that their perception of palatability has not diminished. D) Palatability assessment based on poking structure. Upper pane: poking bout length analysis. A poking bout was defined as 5 consecutive taste deliveries (no withdrawal from nose poke) or more. A bout ends when the rat does not poke for at least 10s. Lower pane: bout length across groups. Distribution of bout length was similar across BLA_ox_ groups and control. *** p<0.001

Another possibility is that BLA_ox_ interrupts with palatability perception. Sucrose is a highly palatable taste and rats drink it avidly even when it is novel (Miller and Holzman, 1981). We reasoned that if the sucrose becomes less palatable with BLA_ox_, we should observe a reduction of sucrose consumption in the experimental group compared with the control in the Pre-CTA training day. Our results, however, show that all BLA_ox_ groups similarly increased their consumption of sucrose compared with the previous day (Fig. 3C, Two-way ANOVA, Taste: F(1,5)=68.4, p=2.0×10^-12^; Group: F(1,5)=1.72, p=0.137, Interaction: F(1,5)=1.37, p=0.241, post-hoc t-tests were all with p<0.001). Another method for palatability assessment uses analysis of licking bouts (Davis, 1989; Hsiao and Fan, 1993; Spector et al., 1998). Using this method with poking bouts (Fig. 3D, upper panel) revealed similarity in bout length between the BLA_ox_ and control groups, indicating no interference of the BLA_ox_ with the perceived high palatability of the sucrose (Fig. 3D lower panel, One-way ANOVA, F_(4)_=1.67, p=0.153). The group with the 5000ms ISI was excluded from this analysis since its bout distribution was incomparable to the 3000ms trials. These results further support our previous assertion regarding the lack of BLA involvement in taste identity perception since licking behavior should depend on taste identity. Together, these results indicate that CTA impairment following BLA inhibition cannot be explained by indirect changes in taste identity or palatability perceptions.

### CTA learning is mediated by LE-BLA activity through the BLA→NBM pathway

Since the basic perceptions of identity and palatability were intact under BLA_ox_, we moved on to test whether BLA-LE activity is essential for the taste novelty processing and the creation of a novel TMT. Novelty processing and TMT are probably distributed across several brain regions (Bermúdez-Rattoni, 2004; Bermudez-Rattoni, 2014), but there is an overall agreement that the GC is one of its primary locations (Bermúdez-Rattoni, 2004; Spector, 2009; Adaikkan and Rosenblum, 2012). The creation of the TMT in the GC is associated with cholinergic signaling from the NBM (Miranda et al., 2000; Power et al., 2002; Power, 2004; Bernstein and Koh, 2007; Rosenberg et al., 2016). Since the BLA projects to the NBM and the BLA→NBM pathway has been shown to enhance other types of memory (Power et al., 2002), we hypothesized that CTA learning is mediated by LE-BLA activity through the BLA→NBM pathway, probably to support novelty processing and TMT formation. To test this hypothesis, we infected the BLA of rats with an ArchT-carrying virus as before (Fig. 4A), but now implanted the fiber optics above the NBM (Fig. 4B, see Materials and Methods) in order to selectively inhibit only BLA afferents into the NBM. Inspecting brain sections following BLA infection revealed dense stained projections from the BLA in the NBM (as well as in other brain areas such as the stria terminalis) (Fig. 4C). We repeated the CTA protocol with LE-BLA_ox_ during the Pre-CTA training session, but now only the BLA projections to the NBM were inhibited (LE-BLA→NBM_ox_, n=4) for a brief 500ms between 700-1200ms following each taste stimulus. Our results show that LE-BLA→NBM_ox_ significantly attenuated CTA learning compared with controls (n=3) infected with a sham virus (Fig. 4D, t-test p<0.001), and similarly to LE-BLA_ox_ performed within the BLA (shown in Fig. 2D, group 700-1200ms, t-test, p=0.425). This result shows that the influence of LE-BLA activity on CTA acquisition is transmitted through the BLA→NBM pathway, and further supporting the role of LE-BLA activity in the novelty processing and the creation of the novel-TMT.

**Fig 4:**
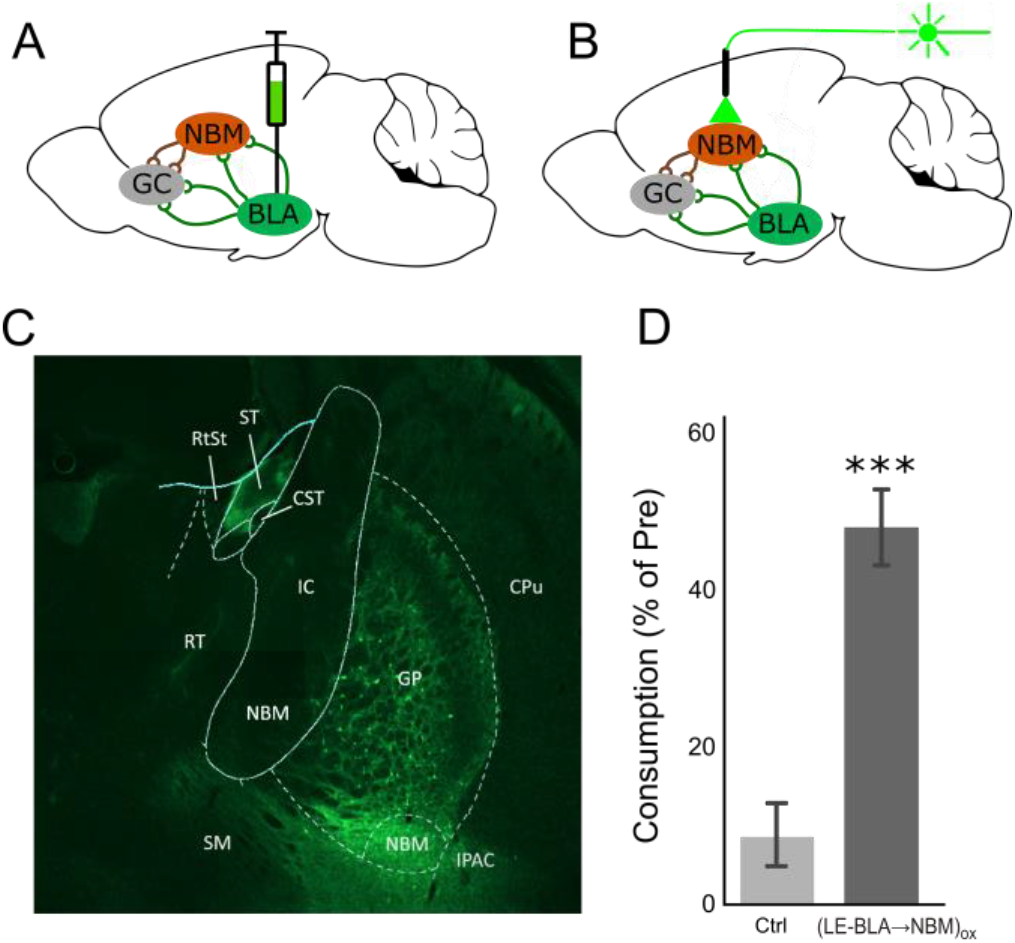
Inhibiting the BLA→NBM pathway attenuates CTA learning. A-B) Schematic illustration of the preparation. A) AAV injection in the BLA followed by B) Fiber optic implantation above the NBM. C) Fluorescent image of infected BLA axon terminals in the NBM area showing dense expression. D) Consumption on the Post-CTA day relative to Pre-CTA day for control and LE-BLA→NBM_ox_ rats. Rats receiving LE-BLA→NBM_ox_ show attenuated CTA learning.

### LE-BLA activity during novel taste experience increases c-Fos expression in the GC

The high levels of NBM-secreted ACh in the GC in response to a novel taste initiates molecular cascades in GC neurons which take part in the formation of the novel-TMT (Berman et al., 2000; Rosenblum et al., 2000; Gutierrez et al., 2003; Rosenberg et al., 2016). Transcription of c-Fos, an immediate early gene, is a fundamental part of this process (Bernstein and Koh, 2007). If LE-BLA activity promotes (presumably through the NBM) the acquisition of a novel taste memory, then inhibiting this activity should also prevent the increased expression of c-Fos in the GC. To test this, we trained rats to poke for water over 4 days and then replaced the water with a novel sucrose solution. Experimental groups received BLA_ox_ during taste deliveries. Ninety minutes later (during the peak of a novel taste associated c-Fos expression (Koh et al., 2003; Wilkins and Bernstein, 2006; Doron and Rosenblum, 2010; Lin et al., 2012b) brains were harvested and sections from the GC area were cut and immunostained for cell nuclei (DAPI) and c-Fos (see the Methods section). Examples of stained slices imaged using fluorescent microscopy are presented in Fig. 5A. To test the role of LE-BLA activity we counted the number of c-Fos expressing neurons in the GC (Fig. 5B). As previously reported, we found higher number of stained c-Fos cells in rats exposed to a novel sucrose (S) compared to a familiar sucrose (S-fam) (Fig. 5C, t-test, p=0.0006) (Koh et al., 2003; Lin et al., 2012b). In the experimental groups receiving BLA_ox_, rats exposed to either full (S-BLA_ox_) or short LE-BLA_ox_ (S-LE-BLA_ox_) during novel sucrose experience showed significantly lower cFos expression compared with the novel sucrose (S) group (Fig. 5C; t-test, p=4.7×10^-5^ and p=1.1×10^-3^ respectively). Laser-only effects were rejected using two additional control groups that received water as stimulus, one with BLA_ox_ (W-BLA_ox_) and the other without (W), both showing similar low c-Fos expression (t-test, p=0.77). Further, consumption comparison between the 4 sucrose groups (S, S-fam, S-BLA_ox_ and S-LE-BLA_ox_) did not reveal significant differences, thereby confirming that the reduction in the GC c-Fos count in the S-BLA_ox_ and S-LE-BLA_ox_ groups is not the result of the lowered sucrose stimulation (Fig. 5D, One-way ANOVA F_(1,3)_=0.34, p=0.71). These results further support a causal relation between the LE-BLA activity and the activation of molecular pathways creating the TMT in the GC.

**Figure 5:**
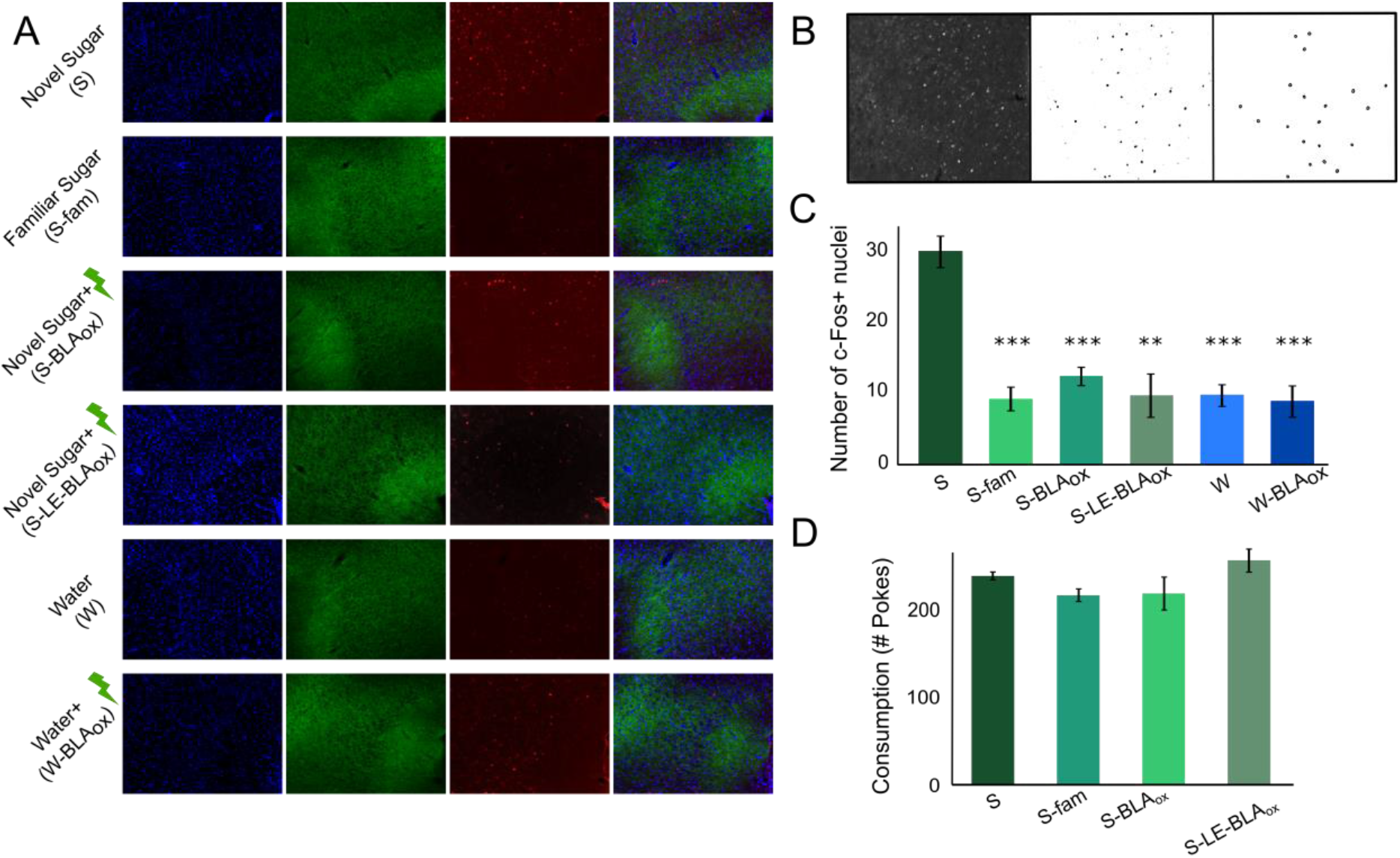
LE-BLA activity during novel taste experience is required for GC c-Fos expression. A) 4’,6-diamidino-2-phenylindole (DAPI) nuclear counterstain staining (in blue) with GFP of BLA projection neurons infected with AAV virus (in green) and c-Fos staining (in red) of the GC in rats from groups receiving different stimulants and BLA_ox_ protocols (scale bar = 200um) B) Example of particle analysis steps used in order to count nuclear colocalizations of DAPI and c-Fos. A threshold was first calculated for each raw image (left) to produce a binary image (center) from which round objects in a specific size range were counted (right) C) Quantification of c-Fos+ positive nuclei in GC neurons. Control rats which drank sucrose without BLA_ox_ (S group) showed a significantly higher expression pattern when compared to rats drinking either a familiar sucrose (S-fam), or rats drinking a novel sucrose with full (S-BLA_ox_) or late-epoch (S-LE-BLA_ox_). Two additional control groups that received water without or with BLA inhibition (W, and W-BLA_ox_, respectively) rejects laser-only effects. D) Average sucrose consumption during Pre-CTA was similar between groups and control, thus cannot account for the differences in c-Fos expression patterns are not due to changes in drinking patterns. *** p<0.001, ** p<0.01.

### LE-BLA activity is critical for valence coding changes in GC following CTA

Changes in behavior following learning are the consequence of changes in neuronal firing activity. Following CTA, single-neuron taste responses (Yasoshima et al., 1995; Moran and Katz, 2014) and ensemble dynamics (Moran and Katz, 2014) in the GC change with transitions in the palatability valence (from palatable to aversive). We therefore tested whether the observed attenuated behavior following LE-BLA_ox_ is the result of disrupted palatability coding in the GC. To do this, we infected the BLAs of rats as before to allow BLA_ox_ (Fig. 6A top), but in addition to the optic fibers we implanted a bundle of electrodes in the GC (Fig. 6A bottom). Rats underwent the same CTA procedure as before, but each daily session was divided into two parts: an initial 20 minutes of free poking for drops of liquid delivered through the IOC with 3 seconds ISI (Fig. 6B, top), followed by forced delivery of taste drops through the IOC with 20 seconds ISI (Fig. 6B, bottom) to record neuronal taste responses. A control group (Ctrl) was infected with sham virus and passed the same procedure as the experimental group. We recorded in total 184 neurons (Ctrl-Pre [n=71], Ctrl-Post [n=43], BLA_ox_-Pre [n=24] and BLA_ox_-Post [n=46]) from 4 control and 5 BLA_ox_ rats. When we tested the effect of random short BLA_ox_ on the baseline activity of GC neurons, we found that it elicited either excitation (Fig.6C, left) or inhibition (Fig. 6C, right), similarly to what was found using long term *in-vivo* BLA pharmacological inhibition (Piette et al., 2012), and in agreement with *in-vitro* studies of BLA→GC connectivity (Haley et al., 2016). While more neurons were inhibited than excited (Fig. 6D), the increased firing rate in excited neurons was about 2 times larger than the decrease in inhibited neurons (59.3% vs. 28.4%, respectively). As expected, the ratio between sampled excitatory and inhibitory neurons were similar between the groups (Fig. 6E) (χ^2^ test, p=0.63). In addition, no difference was observed between the groups in the population averaged baseline and taste-evoked firing across days (Fig. 6F).

**Figure 6:**
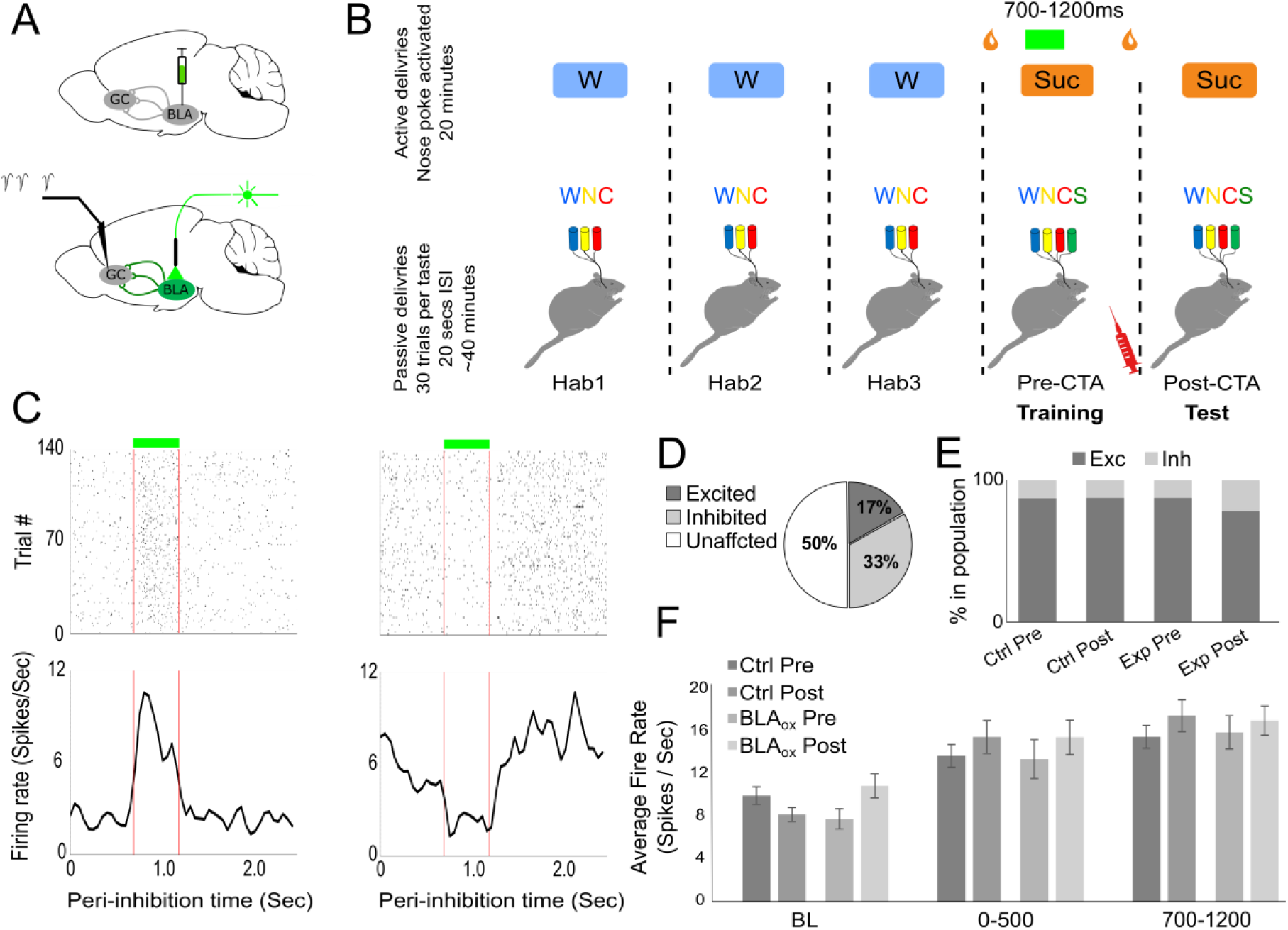
Experimental paradigm for testing palatability coding changes in GC neurons following CTA with LE-BLA_ox_. A) Top: Rats were first bilaterally infected with viruses containing either ArchT or GFP (controls) in the BLA. Bottom: Following recovery simultaneous LE-BLA optogenetic inhibition and GC neuronal recording were performed. B) Experimental procedure: Each daily session was divided into two parts: initial 20 minutes of active poking for liquids (top) followed by forced tastes deliveries for taste response recording (W – water, N – NaCl, C/CA – citric acid, S – sucrose). Rats received LE-BLA_ox_ (Green bar) during the poking session of the training day. Injection of LiCl was given at the end of the forced taste delivery session. C) Example GC neurons being either excited (left) or inhibited (right) by 0.5 second BLA_ox_. Top: Raster plot of spikes. Bottom: Average firing rate over all trials, presented as peri-inhibition time histogram. Red lines delimit BLA_ox_ time. D) Percentage of neurons that were excited, inhibited or unaffected by BLA_ox_ E) Percentage of putative excitatory and inhibitory neurons in the recorded GC neuronal population. F) Mean firing rate of neuron before (BL), 0-500ms after and 700-1200ms after taste delivery from BLA_ox_ and control animals shows no significant difference between groups and group x day interaction. Two-way ANOVA: BL Group p=0.8, Day p=0.99, Group x Day p=0.28; 0-500 Group p=0.92, Day p=0.44, Group x Day p=0.95; 700-1200 Group p=0.9, Day p=0.46, Group x Day p=0.68

We proceeded to test the impact of LE-BLA activity on the post-CTA update of GC palatability coding. First, the spiking activity in response to each taste delivery was used to calculate the post-stimulus time histogram (PSTH, Fig. 7A) for each neuron and each of the tastes. We used these PSTH responses to define the “Palatability Distance to sucrose” (PD, see Materials and Methods). PD is high (above 1) when the response to sucrose is typified by firing dynamics that are similar to the response to the palatable salt solution, and low (below 1) when this response is similar to aversive citric acid.

**Figure 7:**
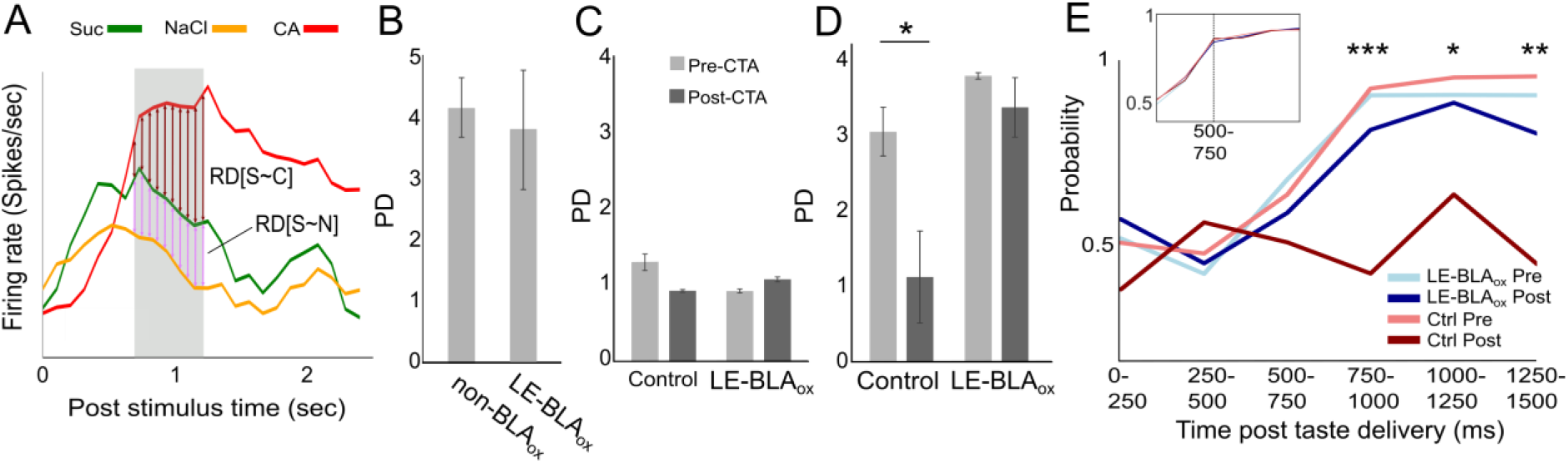
LE-BLA activity during taste exposure is important for CTA-related GC neurons palatability update. A) Palatability distance (PD) ratio calculation for a representative neuron. Colored lines represent mean responses of the neuron to sucrose (Suc), NaCl and citric acid (CA) solutions. Vertical colored arrows in the 700-1200 ms epoch (shaded grey area) mark the distances used for the calculation of response differences (RD), brown arrows for the RD[S~C], and pink for RD[S~N]. The ration between the RDs was used for PD calculation. B) The effect of LE-BLA_ox_ on palatability coding during training. Pre-CTA PD values during the active poking (with BLA_ox_), and from the passive deliveries (non-BLA_ox_) during the LE period in the experimental group. C, D) The effect of LE-BLA_ox_ on palatability coding following CTA. C) PD of sucrose during the EE (0-500ms) was low for all groups before and after CTA. D) LE (700-1200ms) sucrose palatability was high in both groups before CTA. While CTA caused a significant decrease of palatability in the control group, it remained high and similar to Pre-CTA levels in the LE-BLA_ox_ group. E) Population-level palatability coding changes following CTA but not under LE-BLA_ox_. Time-binned GC neuronal responses to sucrose were classified as either NaCl (palatable) or CA (aversive) using a classifier trained with only NaCl and CA trials. Pre-CTA sucrose response of control and LE-BLA_ox_ groups were equally successful in classifying sucrose trials as palatable in the LE bins (starting at 750ms). Post-CTA sucrose trials of the control group were significantly less likely to be classified as palatable NaCl, while those of the LE-BLA_ox_ remained high, similar to Pre-CTA levels. Two-way ANOVA; Group: F(3)=8.2 p=2.2×10^-5^, Bin: F(6)=20.5 p=1.8×10^-21^, Interaction: F(18)=3.2 p=9.9×10^-6^. T-tests between control and LE-BLA_ox_ groups in Post-CTA sessions: Bin0-250 p=0.09, Bin250-500 p=0.48, Bin500-750 p=0.54, Bin750-1000 p=0.0006, Bin100-1250 p=0.01, Bin1250-1500 p=0.002. Inset: The classifier’s performance in correctly identifying NaCl and CA trials, reaching ~90% success 500ms after taste stimulation.

We first wanted to test whether LE-BLA_ox_ during training (before CTA) affects palatability coding in GC. In figures 3C and 3D we showed that BLA_ox_, including LE-BLA_ox_, does not change the palatability of sucrose (high consumption of sucrose with BLA inhibition). This lack of BLA_ox_ impact over palatability was also apparent in GC activity profiles: Similar and high PD values where found when sucrose trials were examined either during the active poking period (with LE-BLA_ox_) or during the passive deliveries (non-BLA_ox_) (Fig. 7B, paired t-test, p=0.88). These results suggest that LE-BLA activity is not essential for GC representations of palatability information.

Next, we studied the importance of LE-BLA epochal activity on the post-CTA update of palatability coding in the GC. This was done by comparing Pre-to Post-CTA PD values calculated from the forced taste deliveries in the second part of the sessions (see Fig. 6B, bottom). As expected from previous studies (Katz et al., 2001; Piette et al., 2012; Sadacca et al., 2012), GC-EE palatability information in both control and LE-BLA_ox_ groups showed low palatability content (Fig. 7C). Correspondingly, PD values were similarly unchanged by CTA learning in both groups (Two-way ANOVA, Day: F_(1,1)_=0.61, p=0.43; BLA_ox_: F_(1,1)_=0.64, p=0.42; Interaction: F_(1,1)_=3.46, p=0.065) (Fig. 7C). In contrast, the palatability-rich GC-LE indeed showed a 3-fold increase in palatability relative to the EE in both groups (Fig. 7D). This increase was similar for both groups (t-test, p=0.13). The groups, however, differ in their update of palatability following CTA training: while there was a significant decrease in palatability in the control group (t-test, p=0.04), palatability in the LE-BLA_ox_ group remained high and similar to its Pre-conditioning values (t-test, p=0.367). These single-neuron level results are in striking agreement with the rats’ behavior across days: rejection of sucrose by the control group and normal consumption by the LE-BLA_ox_ group (Fig. 2E).

Previous work also showed that CTA changes population-level palatability representation in GC neurons (Grossman et al., 2008; Moran and Katz, 2014). We therefore wanted to test whether LE-BLA activity also influences the update of this representation. To test this we aggregated neurons from all rats into 4 groups according to the experimental day and condition (Crtl-Pre, Ctrl-Post, LE-BLA_ox_-Pre, LE-BLA_ox_-Post). We then trained multilayered classifiers to distinguish between salt (palatable) and acid (aversive) solutions using the neuronal responses (recorded in the passive delivery part of the sessions, see Materials and Methods). A separate classifier was built for each of the six 250ms time bins, 0-1500ms post-taste. This allowed us to examine population-coded sucrose palatability across time. The performance of these classifiers in identifying the salty and acid solution showed near chance-levels during the early activity (0-250ms) and increased to ~90% correct classification from 500ms onwards (Fig. 7E, inset), indicating that taste identity can be read out from early GC population responses.

We used these classifiers to evaluate the role of LE-BLA_ox_ in the update of palatability information in GC neurons following CTA. To do that, we used the trained classifiers to classify the population responses to sucrose as either NaCl (palatable) or CA (aversive). Prior to CTA, both the control and experimental groups start at chance level in the EE and increase to ~90% during LE. These results indicate that on the population-level, sucrose is classified as palatable. Following CTA the control group significantly lowered its classification of sucrose as palatable (Fig. 7E), in accord with the behavioral results showing lower sucrose consumption (Fig. 2E). In contrast, in the experimental group that received LE-BLA_ox_, most trials were still classified as palatable. This analysis, therefore, clearly shows that the LE-BLA_ox_ caused a lack of update in sucrose palatability coding that remains “palatable” even after CTA training. Our electrophysiological results indicate that the LE-BLA activity is a critical prerequisite for CTA learning that affects the update of GC neurons’ palatability coding, in both the single and ensemble levels.

## Discussion

Here we examined the roles BLA activity plays during taste perception and memory in the taste system, and the pathways by which this activity is transmitted. Collectively, our experiments suggest a distinct role of LE activity of BLA projection neurons (~700-3000 ms from initial taste experience) in novelty processing and the formation of a novel-taste memory trace in the GC. We show that BLA_ox_, and specifically the LE-BLA_ox_, interfere with the acquisition of CTA memory (Fig. 2), without disruption of taste identity or palatability perception (Fig. 3). In addition, we show that LE-BLA activity during novel taste stimulation is required for c-Fos expression in the GC (Fig 5) – a known marker of the novel-taste memory formation in the GC. Correspondingly, LE-BLA activity during novel taste experience is required for the update of palatability coding in GC following CTA (Fig 7). Lastly, we were able to show that the BLA→NBM pathway is an essential pathway for transmission of LE-BLA information required for CTA (Fig 4). Together, these results reveal a circuit for novel taste processing that promotes the creation of taste recognition memory across the BLA, NBM and GC through temporally specific neuronal dynamics.

The BLA has been implicated in novel taste processing for a long time (Nachman and Ashe, 1974; Miranda et al., 2003b; Reilly and Bornovalova, 2005; St Andre and Reilly, 2007). However, there is no clear system-level understanding of how novelty information propagates across regions. The only electrophysiological study that investigated familiarization coding was done in the GC and showed a correlation between familiarity and neuronal firing rate, but only in a later phase of the response (>2 sec) (Bahar et al., 2004). Using the same logic as for palatability coding (which is detected earlier in the BLA than in GC), our results support the hypothesis that novelty information is first processed in the BLA and then sent to the GC. Whether the BLA itself is the source for taste novelty or familiarity, or whether it receives this information from different brain areas remains an open question. Answering this question will require a systematic search for novelty/familiarity correlates across the taste system.

One of the main contributions of this study is to show the distinct roles played by the BLA in shaping taste-related behaviors. Short-timed LE-BLA_ox_ attenuated the novelty-related learning of CTA, however even a complete BLA inhibition showed no effect on the ability of the rats to identify a taste (Fig. 3B) or to perceive its palatability (Fig. 3C, D). The lack of BLA_ox_ impact on perceptual-sensitive tasks seems inconsistent with studies showing correlations between neuronal activity in the BLA and palatability processing (Fontanini et al., 2009) and reduced GC palatability information following BLA pharmacological inhibition (Piette et al., 2012). At least two propositions may reconcile these inconsistencies. According to the first, the BLA is indeed involved in taste palatability perception, but it is not exclusive in doing so. Correspondingly, other brain areas were shown to be involved in palatability processing, and may compensate for the lack of BLA input such as the parabrachial pontine nucleus (PbN) (Baez-Santiago et al., 2016), medial prefrontal cortex (Jezzini et al., 2013) and the nucleus accumbens (Taha and Fields, 2005). The second option is that although there exist a correlation between neuronal firing rate and palatability, the role of neuronal activity in BLA and GC is not primarily to support perception but rather to promote memory and learning processes, such as the formation of a novel-TMT and CTA (Fig. 3C, D). In accord with both of these interpretations, several studies reported uninterrupted consumption following BLA lesion or pharmacological inhibition during the training day of the CTA procedure, before any learning (Rolls and Rolls, 1973; Nachman and Ashe, 1974; Gallo et al., 1992). Therefore, our study calls for a careful interpretation of correlations between neuronal activities and assumed perceptions without an additional functional proof of their causal relations.

It is important to note that to assess taste perception and memory formation while disregarding neophobia requires the usage of a taste that minimizes the neophobic reaction. This was our reason for choosing sucrose, which evokes only minor neophobic responses (Miller and Holzman, 1981; Franchina and Gilley, 1986; Franchina and Slank, 1988; Flores et al., 2016), as opposed to saccharin (Domjan and Gillan, 1976; Lin et al., 2012a). This choice, however, prevented us from exploring the relation between BLA epochal activity and neophobic reaction. A recent study which used BLA pharmacological inhibition showed that the BLA is essential for the occurrence of neophobia (Lin et al., 2018). These results are inline with our results, emphasizing the role of the BLA in novelty-sensitive behaviors. Whether the same BLA epoch is responsible for both the neophobic reaction and the formation of the TMT remains to be discovered.

The BLA may influence the GC during novel taste experience directly and/or indirectly through its connectivity with many other brain regions. In our work, we specifically studied the role of the BLA→NBM pathway (Fig. 4D). A novel taste is known to evoke a novelty signal of extracellular ACh secreted from NBM→GC afferents (Miranda et al., 2000, 2003a; Bermúdez-Rattoni, 2004). The elevated ACh in the GC triggers a complex molecular cascade that includes, among other, ERK-I/II phosphorylation (Rosenblum et al., 2000), c-Fos activation (Clark and Bernstein, 2009), and decrease in proteasome activity (Rosenberg et al., 2016), which is known to underlie the creation of the novel-TMT (Bermúdez-Rattoni, 2004). The BLA is a valid candidate for a NBM activator following the introduction of a novel taste. Anatomically, the BLA densely innervates the NBM (Grove, 1988; Jolkkonen et al., 2002) and can potentially influence its activity. Functionally, the BLA→NBM and NBM→cortical cholinergic projections are known to be important players in the modulation and formation of memories in other modalities (Dringenberg and Vanderwolf, 1996; Power et al., 2000; McGaugh, 2002). Accordingly, we showed that the BLA→NBM pathway, and specifically the LE activity of this pathway, is vital for the creation of CTA. Based on the collective results obtained here and in previous studies (Jolkkonen et al., 2002; McGaugh, 2002; Miranda et al., 2003a; Bermúdez-Rattoni et al., 2004) we suggest that the BLA→NBM pathway participates in novelty processing and the formation of the novel-TMT in the GC, that later results in the CTA memory formation. We note, however, that we do not claim that the novel-TMT is strictly confined to the GC. The engram of the novel-TMT is probably distributed across different brain regions, even in the BLA itself, as suggested by the increase in c-Fos expression in the BLA following novel taste consumption (Lin et al., 2012b). It was also shown that blocking ACh input to the GC alone is not sufficient for attenuating CTA, but reducing ACh input to both GC and BLA is (Gutiérrez et al., 1999). Similarly, noradrenergic receptor blocking in the BLA following novel taste experience attenuate familiarity learning (Miranda et al., 2003b). These results suggest that changes in BLA are also important for novelty processing and the creation of the novel-TMT.

In summary, our results demonstrate the importance of the LE-BLA during novel taste experiencing to the formation of taste memories in the GC. The formation of these memories crucially depends on the currently understudied BLA→NBM pathway. We suggest that LE-BLA activity during novel taste experiencing is important for processing taste novelty, the formation of a TMT in the GC, and following malaise, to the creation of CTA memory. Our results highlight the importance of neuronal dynamics in sensory information processing and in inter-regional communication, memory formation and learning.

## Acknowledgments

We thank Daniel Udi, Danit Miron and Lihi Cohen for technical assistance. We thank Prof. Kobi Rosenblum, Dr. David Levitan and Dr. Veronica Flores for guidance regarding the c Fos staining. The research was supported by seed grant from the Tel Aviv University.

